# A genetically encoded GRAB sensor for measuring serotonin dynamics *in vivo*

**DOI:** 10.1101/2020.02.24.962282

**Authors:** Jinxia Wan, Wanling Peng, Xuelin Li, Tongrui Qian, Kun Song, Jianzhi Zeng, Fei Deng, Suyu Hao, Jiesi Feng, Peng Zhang, Yajun Zhang, Jing Zou, Sunlei Pan, J. Julius Zhu, Miao Jing, Min Xu, Yulong Li

## Abstract

Serotonin (5-HT) is a phylogenetically conserved monoamine neurotransmitter modulating a variety of processes in the brain. To directly visualize the dynamics of 5-HT, we developed a genetically encoded <u>G</u>PC<u>R</u>-<u>A</u>ctivation-<u>B</u>ased 5-HT (GRAB_5-HT_) sensor with high sensitivity, selectivity, and spatiotemporal resolution. GRAB_5-HT_, detected 5-HT release in multiple physiological and pathological conditions in both flies and mice, and thus provides new insights into the dynamics and mechanisms of 5-HT signaling.

## Main

Serotonergic signaling in the brain plays a critical role in a wide range of physiological processes, including mood control, reward processing, and sleep-wake homeostatic regulation^1–3^. Given its functional importance, drugs targeting central serotonergic activity have been used to treat virtually every psychiatric disorder, with the best example being the use of selective serotonin reuptake inhibitors (SSRIs) for depression^4^. Despite the importance of 5-HT, understanding of cell-specific 5-HT signaling during behaviors is greatly hampered by the lack of ability to measure 5-HT *in vivo* with high sensitivity and precise spatiotemporal resolution^5,6,7^. Using molecular engineering, we developed a genetically encoded fluorescent sensor for directly measuring extracellular 5-HT.

Previously, we and others independently developed GPCR activation based sensors for detecting different neurotransmitters by converting the conformational change in the respective GPCR to a sensitive fluorescence change in circular permutated GFP (cpGFP)^8,9,10,11^. Using similar strategy, we initiated the engineering of 5-HT‒specific GRAB sensor by inserting a cpGFP into the third intracellular loop (ICL3) of various 5-HT receptors. Based on the performance of membrane trafficking and affinity of receptor-cpGFP chimeras, we selected and focused on the 5-HT_2C_ receptor‒based chimera for further optimization (Extended Data Fig. 1a,b). Mutagenesis and screening in its linker regions and cpGFP moiety, resulted in a sensor with a 250% ΔF/F_0_ in response to 5-HT, which we named GRAB_5-HT1.0_ (referred to hereafter as simply 5-HT1.0; Fig 1a and Extended Data Fig. 2). In addition, we generated a 5-HT‒insensitive version of this sensor by introducing the D134^3.32^Q mutation in the receptor^12^, resulting in GRAB_5-HTmut_ (referred to hereafter as 5-HTmut). This mutant sensor has the similar membrane trafficking as 5-HT1.0, but <2% ΔF/F_0_ even in response to 100 μM 5-HT (Fig. 1a and Extended Data Fig. 3a-d). In cultured neurons, the 5-HT1.0 sensor produced a robust fluorescence increase (280% ΔF/F_0_) in both the soma and neurites in response to 5-HT application, whereas 5-HTmut sensor had no measurable change in fluorescence (Fig. 1b and Extended Data Fig. 3l).

**Fig1:**
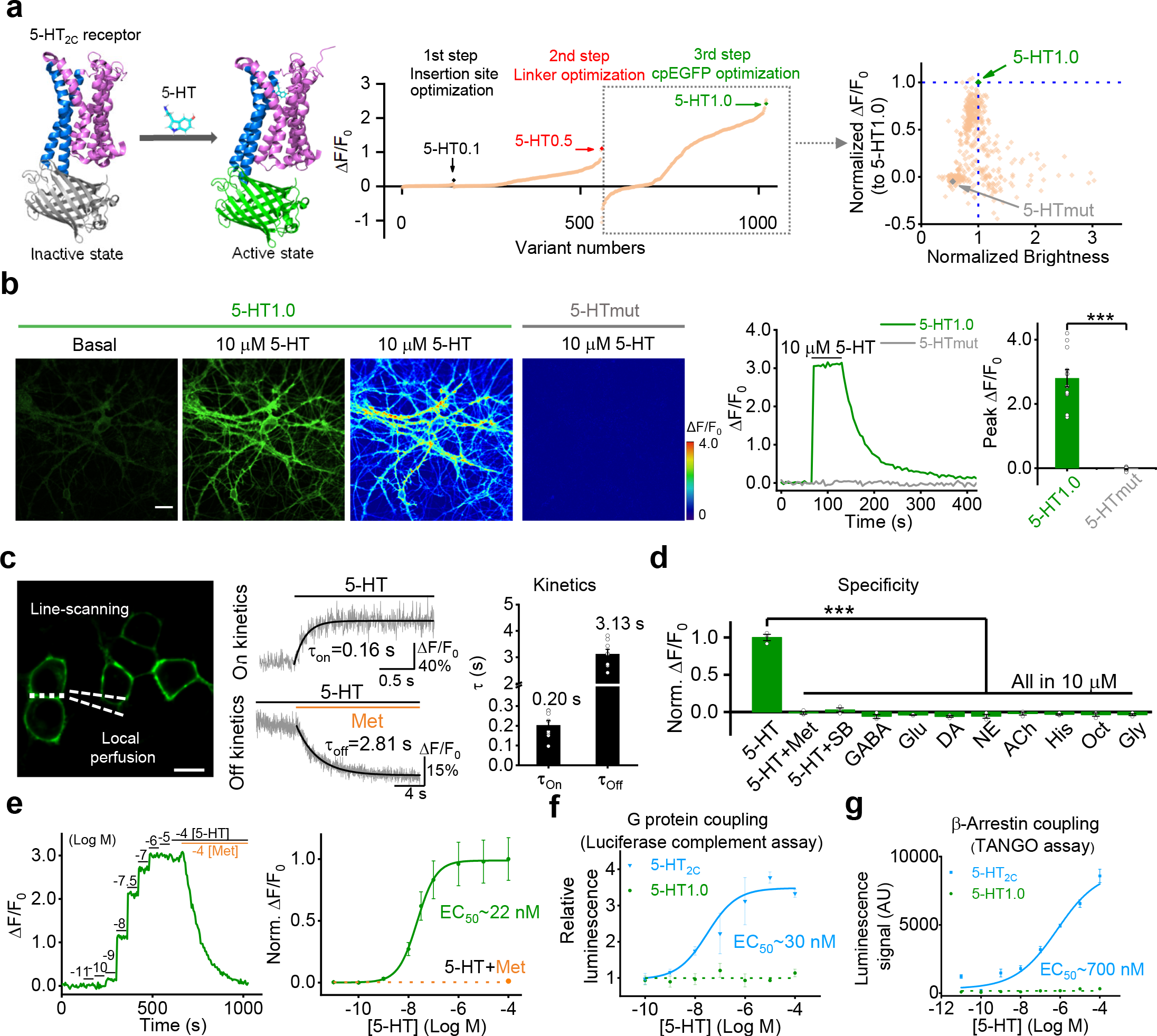
Design, optimization, and characterization of a novel genetically encoded 5-HT sensor. **(a)** Left: schematic representation illustrating the principle behind the GRAB5-HT sensor. The crystal structures are from Protein Data Bank (PDB) archive (PDB ID: 6BQH and 6BQG for the inactive and active states of the 5-HT_2C_ receptor, respectively^12^, and PDB ID: 3EVP for cpGFP^31^). Middle: the 5-HT sensor was optimized over 3 main steps, including the cpGFP insertion site, the linker between cpGFP and 5-HT_2C_, and critical amino acids in cpGFP. Right: optimization of cpGFP and the engineering of 5-HTmut. The fluorescence change in each candidate sensor is plotted against the brightness, with both axes normalized to 5-HT1.0. **(b)** Representative images (left), fluorescence traces (middle), and group data (right) of the fluorescence response in neurons expressing 5-HT1.0 (green) or 5-HTmut (gray); where indicated, 10 μM 5-HT was applied; n = 12/3 (12 cells from 3 cultures) for each group. Scale bar, 20 μm. **(c)** Kinetic analysis of the 5-HT1.0 sensor. Left, a representative image showing the experiment protocol, in which the line-scanning mode was used to record the fluorescence change in cells expressing 5-HT1.0 in response to local application of 5-HT, followed by metergoline (Met) in the continued presence of 5-HT. Middle, representative traces showing the rise and decay of 5-HT1.0 fluorescence in response to 5-HT (top) followed by Met (bottom). Right, a summary of on and off kinetics of 5-HT1.0; n = 8/5 for each group. Scale bar, 10 μm. **(d)** Summary of the change in fluorescence of 5-HT1.0 in response to 5-HT alone, 5-HT together with Met or SB 252084 (SB), and 8 additional neurotransmitters and neuromodulators. GABA, gamma-aminobutyric acid; Glu, glutamate; DA, dopamine; NE, norepinephrine; ACh, acetylcholine; His, histamine; Oct, octopamine; and Gly, glycine; n = 3 wells per group with 300–500 cells per well. **(e)** Dose-response curve measured in neurons expressing 5-HT1.0 in response to increasing concentrations of 5-HT, followed by Met; n = 18/4 for each group. **(f, g)** G-protein coupling **(f)** and β-arrestin coupling **(g)** was measured for the 5-HT1.0 sensor and 5-HT2C receptor using a luciferase complementation assay and TANGO assay, respectively; n = 3 wells per group with 100-300 cells per well. In this and subsequent figures, summary data are shown as the mean ± SEM. *, *p*<0.05, **, *p*<0.01, ***, *p*<0.001, and n.s., not significant (Student’s *t*-test).

Next, we characterized the properties of the 5-HT1.0 sensor, including the brightness and photostability, dose-response relationship between 5-HT concentration and fluorescence change, response kinetics, signal specificity and downstream coupling. We found that the 5-HT1.0 had similar brightness and better photostability compared to EGFP (Extended Data Fig. 3e-g). In addition, we measured the sensor’s kinetics upon application of 5-HT (to measure the on-rate) followed by the 5-HT receptor antagonist metergoline (Met, to measure the off-rate) and measured τ_on_ and τ_off_ values to be 0.2 s and 3.1 s, respectively (Fig. 1c). The 5-HT1.0 was highly sensitive to 5-HT, with an EC_50_ of 22 nM (Fig. 1e). Importantly, none of other neurotransmitters and neuromodulators tested elicited a detectable fluorescence change, and the 5-HT-induced signal was eliminated by the 5-HT receptor antagonist SB 242084 (SB) (Fig. 1d and Extended Data Fig. 3m,n), indicting a high specificity to 5-HT. Unlike the native 5-HT_2C_ receptor, which couples to the intracellular G-protein and β-arrestin signaling pathways, the 5-HT1.0 sensor showed no detectable coupling to either of these two pathways measured by the calcium imaging, G-protein‒ dependent luciferase complementation assay^13^, TANGO assay^14^, and long-term measurements of membrane fluorescence in the presence of 5-HT (Fig. 1f,g and Extended Data Fig. 3h-k).

Next, we measured the dynamics of endogenous 5-HT upon neuronal activation. We expressed either 5-HT1.0 or 5-HTmut in the mouse dorsal raphe nucleus (DRN) by AAV, and prepared acute brain slices three weeks after infection (Fig. 2a). In DRN slices expressing 5-HT1.0, a single electrical pulse evoked detectable fluorescence increases, and the response progressively enhanced with the increase in pulse number or frequency (Fig. 2b,c and Extended Data Fig. 4a). The stimulation evoked-response was repeatable for up to 25 min (Extended Data Fig. 4b) and blocked by the 5-HT receptor antagonist Met, but not the dopamine receptor antagonist haloperidol (Halo; Fig. 2d and Extended Data Fig. 4c,d). In contrast, the same electrical stimuli had no effect on fluorescence in slices expressing the 5-HTmut sensor (Fig. 2b,d). We also measured the kinetics of the fluorescence change in response to 100 ms electrical stimulation and found τ_on_ and τ_off_ values of 0.15 s and 7.22 s (Fig. 2e). We further compared the 5-HT1.0 sensor with existing fast-scan cyclic voltammetry (FSCV) in recording 5-HT by simultaneously conducting fluorescence imaging and electrochemical recording in DRN slices (Fig. 2f). Both methods could sensitively detect single pulse-evoked 5-HT signal and the increase of response following incremental frequencies (Fig. 2g and Extended Data Fig. 4e,f). Importantly, the 5-HT1.0 showed better signal to noise ratio (SNR) compared with FSCV (Fig. 2g).

**Fig2:**
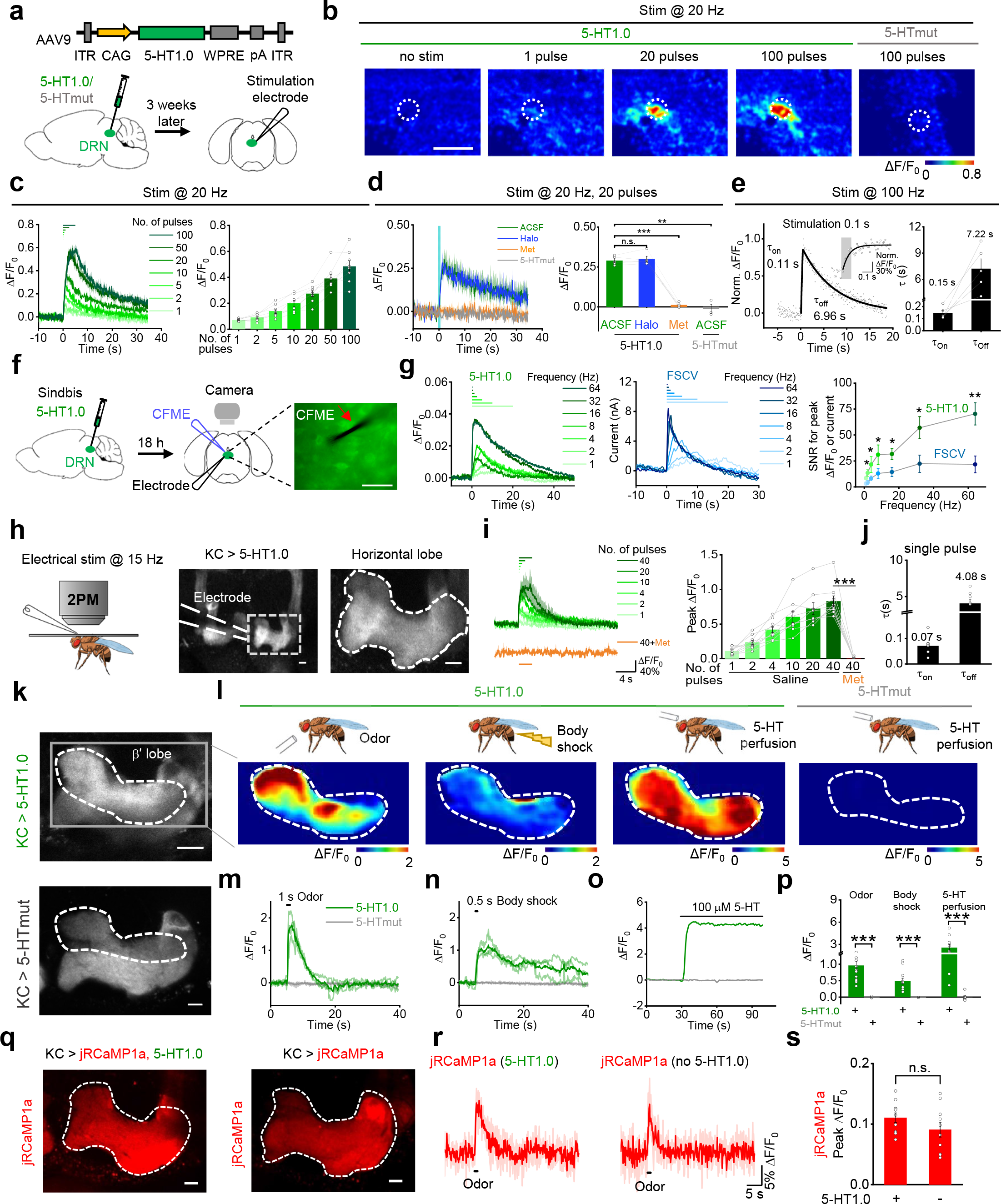
GRAB_5-HT_ can report the release of endogenous 5-HT in acute mouse brain slices and *Drosophila*. **(a)** Schematic illustration depicting the mouse brain slice experiments. Top: The AAV vector used to express the 5-HT1.0 sensor. Bottom: AAV expressing either 5-HT1.0 or 5-HTmut was injected in the mouse DRN, after which acute brain slices were prepared and recorded. **(b)** Representative pseudocolor images of the change in 5-HT1.0 and 5-HTmut fluorescence in response to the indicated electrical stimuli delivered at 20 Hz. The duration of one pulse is 1 ms. The white dotted circle (50 μm diameter) indicates the ROI used for further analysis. Scale bar, 100 μm. **(c)** Individual traces (left) and quantification (right) of 5-HT1.0 fluorescence change in response to the indicated electrical stimuli delivered at 20 Hz; n = 8 slices from 5 mice. **(d)** Representative traces (left) and group data of 5-HT1.0 and 5-HTmut fluorescence change in response to electrical stimuli in slices treated with the dopamine receptor antagonist haloperidol (Halo) or the 5-HT receptor antagonist Met; n = 3-5 slices from 2-4 mice. **(e)** Left: normalized change in 5-HT1.0 fluorescence in response to 10 electrical stimuli delivered at 100 Hz. The rise and decay phases are fitted with single-exponential functions (black traces). A magnified view of the on kinetics in inset. Right: summary of τ_on_ and τ_off_; n = 5 slices from 4 mice. **(f)** Schematic drawing outlines the design of simultaneous imaging and fast-scan cyclic voltammetry (FSCV) experiments in mouse DRN slice preparation. The red arrow indicates that the carbon-fiber microelectrode (CFME) is placed near the neuron expressing 5-HT1.0. Scale bar, 20 μm. **(g)** Left: representative fluorescence traces of a DRN neuron expressing 5-HT1.0 to electrical stimuli consisting of a train of 20 pulses at varied frequency. Middle: current vs time traces of evoked 5-HT release at varied stimulating frequencies. Right: group summary of the signal to noise ratio (SNR) of 5-HT1.0 and FSCV; n = 11 neurons from 9 mice. **(h)** Left: schematic drawing showing *in vivo* two-photon imaging of a *Drosophila*, with the stimulating electrode positioned near the mushroom body (MB). Middle and right: representative images of a fly expressing 5-HT1.0 in the Kenyon cells (KCs) of the MB; the image at the right is a magnified view of the dashed rectangle. The duration of one pulse is 1 ms. Scale bar, 10 μm. **(i)** Representative traces (left) and group analysis (right) of 5-HT1.0 fluorescence in response to the indicated electrical stimuli in either saline (control) or 10 μM Met; n = 9 flies for each group. **(j)** Summary of 5-HT1.0 τ_on_ and τ_off_ in response to a single electrical stimulation; n = 8 flies for each group. **(k)** Fluorescence images measured in the MB of flies expressing 5-HT1.0 or 5-HTmut; the β’ lobe is indicated. Scale bar, 10 μm. **(l-p)** Representative pseudocolor images **(l)**, fluorescence traces **(m-o)**, and group summary **(p)** of 5-HT1.0 and 5-HTmut in the MB β’ lobe measured in response to a 1-s odor application, a 0.5-s body shock, and application of 100 μM 5-HT; n = 5-14 flies for each group. **(q)** Fluorescence images of jRCaMP1a in fly MB with co-expression of 5-HT1.0 (left) or expressing jRCaMP1a alone (right) in the KCs. Scale bar, 10 μm. **(r, s)** Representative traces **(r)** and group summary **(s)** of Ca^2+^ signal in the MB of flies co-expressing jRCaMP1a and 5-HT1.0 or jRCaMP1a alone; where indicated, a 1-s odorant stimulation was applied; n = 10-11 flies for each group.

We next tested whether the 5-HT1.0 sensor could be used to measure sensory-relevant changes in 5-HT signaling *in vivo*. We used the *Drosophila* model, as serotonergic signaling in the mushroom body (MB) has been implicated in odor-related memory consolidation^15^, in which 5-HT is released from single serotoninergic dorsal paired medial (DPM) neurons that innervate Kenyon cells (KC) in the MB per hemisphere^16, 17^. The 5-HT1.0 reliably reported 5-HT release evoked by electrically stimulating the horizontal lobe of MB, revealing rapid on and off kinetics of 0.07 s and 4.08 s, respectively. Moreover, the signal was blocked by applying Met (Fig. 2h-j and Extended Data Fig. 4g-i). Two distinct physiological stimuli—odor application and body shock—evoked a robust fluorescence increase in the MB β’ lobe of flies (Fig. 2k-p and Extended Data movie 1), consistent with previous studies of Ca^2+^ signaling in the DPM^18^. In contrast, no fluorescence changes detected in flies expressing the 5-HTmut sensor (Fig. 2k-p, Extended Data movie 1). Neither stimulus produced a saturated response of 5-HT1.0 sensor, as application of exogenous 100 μM 5-HT in the same flies elicited much larger responses. Finally, co-expressing the 5-HT1.0 sensor together with red fluorescent Ca^2+^ sensor jRCaMP1a in KC to perform two-color imaging to examine whether the expression of the 5-HT1.0 sensor affects the odorant-evoked Ca^2+^ response. Both 5-HT1.0 and jRCaMP1a produced highly sensitive fluorescence increases in response to the odorant application in the green and red channels, respectively (Fig. 2q-s and Extended Data Fig. 4j-l). Importantly, jRCaMP1a-expressing flies with or without co-expression of 5-HT1.0 had similar odorant-evoked Ca^2+^ signals, suggesting virtually no effect of expressing the 5-HT1.0 sensor on cellular physiology (Fig. 2q-s and Extended Data Fig. 4j-l).

Methylenedioxymethamphetamine (MDMA) is a synthetic addictive compound which could alter mood and perception, and its effects can be partially explained by increasing extracellular 5-HT concentrations in the brain^19^. We examined MDMA’s effect *in vivo* by two-photon imaging in mice expressing the sensor in the prefrontal cortex (PFC) (Fig. 3a). Intraperitoneal (i.p.) injection of MDMA caused a progressive increase in 5-HT1.0 fluorescence, which peaked after 1 hour and then gradually decayed over the following 3 hours (Fig. 3b-d). The time course is comparable with previous reports of MDMA’s effects on both human^20^ and mouse^21^. Meanwhile, MDMA had no effect on fluorescence of 5-HTmut sensor (Fig. 3a-d). These results together suggest that the 5-HT1.0 sensor is suitable for stable, long-term imaging *in vivo*.

**Fig3:**
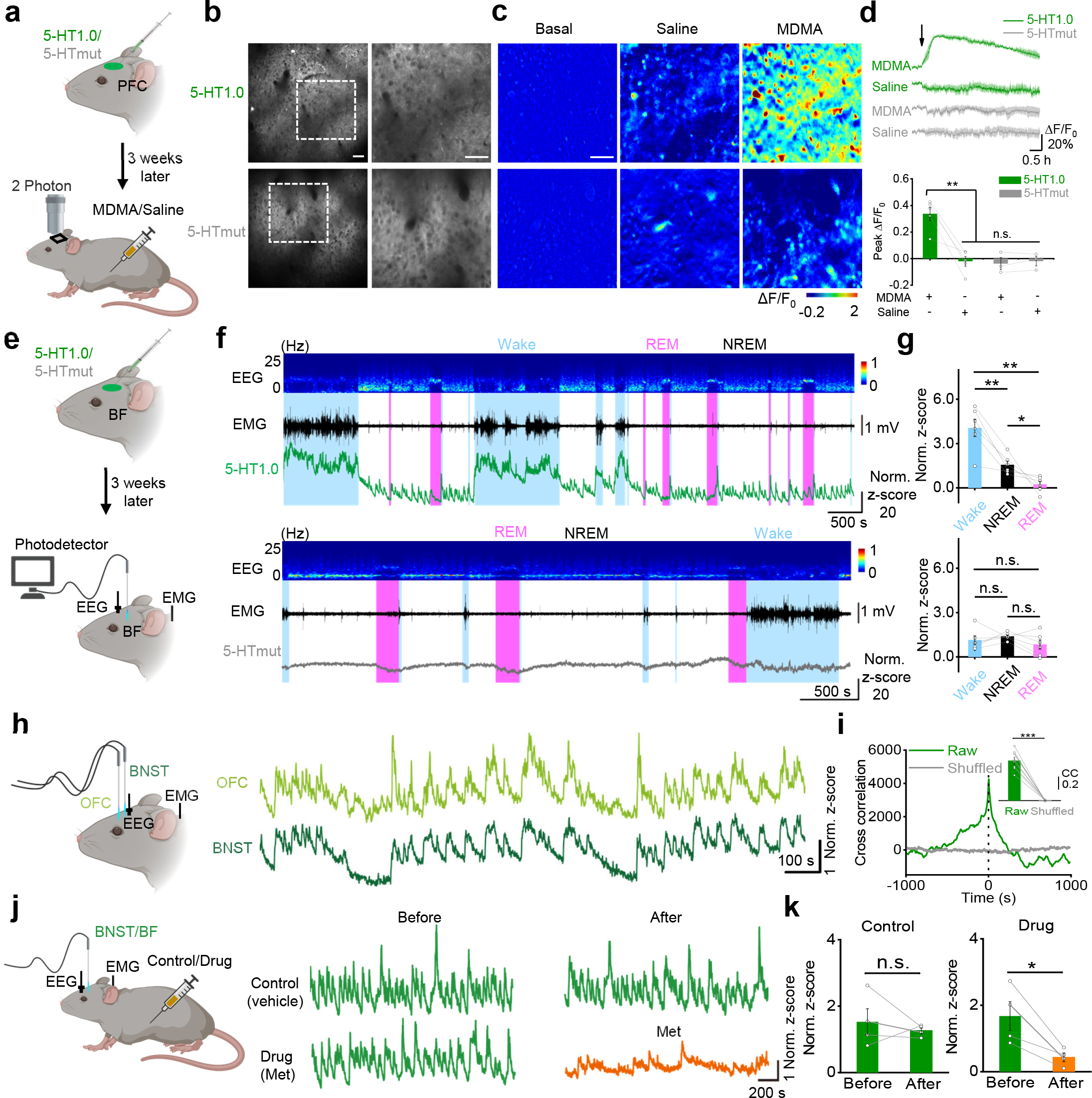
GRAB_5-HT_ can report endogenous serotonergic activity in freely behaving mice. **(a)** Schematic diagram illustrating the use of two-photon imaging to measure 5-HT1.0 and 5-HTmut fluorescence in the prefrontal cortex (PFC) of head-fixed mice; MDMA or saline was injected intraperitoneally. **(b)** Representative images of 5-HT1.0 (top) and 5-HTmut (bottom) fluorescence measured in the mouse PFC. Scale bar, 50 μm. **(c, d)** Representative pseudocolor images **(c)**, averaged fluorescence traces **(d, top)**, and group summary **(d, bottom)** showing 5-HT1.0 (top, green) and 5-HTmut (bottom, gray) fluorescence measured after an i.p. injection of saline (middle) or 10 mg/kg MDMA (right); n = 3-5 mice for each group. Scale bar, 50 μm. **(e)** Schematic diagram illustrating the use of fiber-photometry for measuring 5-HT1.0 and 5-HTmut fluorescence in the basal forebrain (BF) of freely behaving mice during the sleep-wake cycle. EEG and EMG were also measured. **(f)** Representative EEG, EMG, and 5-HT1.0 (top panel) and 5-HTmut (bottom panel) fluorescence measured during the sleep-wake cycle. **(g)** Summary of 5-HT1.0 (top) and 5-HTmut (bottom) fluorescence measured in awake mice and during NREM and REM sleep; n = 3 mice in two sessions for each group. **(h, i)** Same as in **(e)**, except the 5-HT1.0 sensor was expressed in both the orbital frontal cortex (OFC; light green) and the bed nucleus of the stria terminalis (BNST; dark green), and the fluorescence response in each nucleus was recorded and analyzed. The cross-correlation between the signals in the OFC and BNST is shown in **(i)**; n = 4 mice in two sessions for each group. **(j, k)** 5-HT1.0 fluorescence was measured in the BNST and BF as in **(h)**; where indicated, the mice received an injection of saline (control) or Met. The normalized responses in the BNST (n = 3 mice) and BF (n = 1 mouse) were combined for the group summary.

Finally, we examined whether the 5-HT1.0 sensor could measure the dynamics of serotonergic activity under physiological conditions, e.g. the sleep-wake cycle in mice. The 5-HT1.0 sensor was expressed in several brain nuclei, including the basal forebrain (BF), orbital frontal cortex (OFC), and the bed nucleus of the stria terminalis (BNST), then we performed simultaneous fiber-photometry and EEG/EMG recordings in freely behaving mice. In BF, we found that the 5-HT1.0 sensor signal was generally higher when the mice were awake compared to either REM or non-REM (NREM) sleep, with the lowest signal detected during REM sleep, consistent with the notion that 5-HT signaling is minimum during REM sleep^22^ (Fig. 3e-g). As expected, we found no significant change in fluorescence in mice expressing 5-HTmut sensor during the sleep-wake cycle. Interestingly, simultaneous recording 5-HT1.0 in OFC and BNST revealed tight correlation in fluorescence during NREM sleep (Fig. 3h,i), suggesting global synchrony of the 5-HT signaling despite of the region-specific innervation by different subpopulations of the serotonergic neurons in DRN^23, 24^. Lastly, consistent with our previous findings, we found that treating mice with the 5-HT receptor antagonist Met largely blocked the fluorescence change of the 5-HT1.0 sensor (Fig. 3j,k), validating the specificity of measured signals *in vivo*.

In summary, we report the development and application of a novel genetically encoded fluorescent GRAB sensor for measuring extracellular 5-HT dynamics. This GRAB_5-HT1.0_ sensor has high sensitivity and specificity, as well as high spatiotemporal resolution, yet it does not appear to affect cellular physiology. GRAB_5-HT1.0_ reliably reports endogenous 5-HT release in response to a variety of stimuli and under various behaviors in different animal models. GRAB_5-HT1.0_ follows 5-HT dynamics in mice throughout the sleep-wake cycle, providing new insights into the functional contribution of 5-HT in sleep regulation.

## Methods

### Primary cultures

Male and female postnatal day 0 (P0) Sprague-Dawley rat pups were obtained from (Beijing Vital River) and used to prepare cortical neurons. The cortex was dissected, and neurons were dissociated using 0.25% Trypsin-EDTA (GIBCO), plated on 12-mm glass coverslips coated with poly-D-lysine (Sigma-Aldrich), and cultured in neurobasal medium (GIBCO) containing 2% B-27 supplement (GIBCO), 1% GlutaMax (GIBCO), and 1% penicillin-streptomycin (GIBCO). The neurons were cultured at 37°C in a humidified atmosphere in air containing 5% CO_2_.

### Cell lines

HEK293T cells were purchased from ATCC and verified based on their morphology under microscopy and an analysis of their growth curve. Stable cell lines expressing either 5-HT_2C_ or 5-HT1.0 were generated by co-transfecting cells with the pPiggyBac plasmid carrying the target genes together with Tn5 transposase into a stable HEK293T cell line. Cells expressing the target genes were selected using 2 μg/ml Puromycin (Sigma). An HEK293 cell line stably expressing a tTA-dependent luciferase reporter and the β-arrestin2-TEV fusion gene used in the TANGO assay was a generous gift from Bryan L. Roth^25^. All cell lines were cultured at 37°C in 5% CO_2_ in DMEM (GIBCO) supplemented with 10% (v/v) fetal bovine serum (GIBCO) and 1% penicillin-streptomycin (GIBCO).

### Drosophila

UAS-GRAB_5-HT1.0_ (attp40, UAS-GRAB_5-HT1.0_/CyO) and UAS-GRAB_5-HTmut_ (attp40, UAS-GRAB_5-HTmut_/CyO) flies were generated in this study. The coding sequences of GRAB_5-HT1.0_ or GRAB_5-HTmut_ were inserted into pJFRC28^26^ (Addgene plasmid #36431) using Gibson assembly. These vectors were injected into embryos and integrated into attp40 via PhiC31 by the Core Facility of Drosophila Resource and Technology, Shanghai Institute of Biochemistry and Cell Biology, Chinese Academy of Sciences. The following fly stocks were used in this study: R13F02-Gal4 (BDSC:49032) and UAS-jRCaMP1a (BDSC: 63792)^27^. Flies were raised on standard cornmeal-yeast medium at 25°C, with 70% relative humidity and a 12h/12h light/dark cycle. In Fig. 2h-p and Extended Data Fig.4g-i, fly UAS-GRAB_5-HT1.0_/CyO; R13F02-Gal4/TM2 and fly UAS-GRAB_5-HTmut_/+; R13F02-Gal4/+ were used; in Fig. 2q-s and Extended Data Fig.4j-l, fly UAS-GRAB_5-HT1.0_/+; R13F02-Gal4/UAS-jRCaMP1a/+ and fly R13F02-Gal4/UAS-jRCaMP1a were used.

### Mouse

Wild-type C57BL/6 (P25-60) mice were used to prepare the acute brain slices and for the *in vivo* mouse experiments. All mice were group-housed in a temperature-controlled room with a 12h/12h light/dark cycle, with food and water provided ad libitum. All procedures for animal surgery and maintenance were performed using protocols that were approved by the Animal Care & Use Committees at Peking University, the Chinese Academy of Sciences, University of Virginia, and were performed in accordance with the guidelines established by the US National Institutes of Health.

### Molecular biology

Plasmids were generated using the Gibson assembly method^28^, and DNA fragments were amplified by PCR using primers (Thermo Fisher Scientific) with 25-30 bp overlap. DNA fragments were assembled using T5-exonuclease (New England Biolabs), Phusion DNA polymerase (Thermo Fisher Scientific), and Taq ligase (iCloning). Sanger sequencing was performed at the Sequencing Platform in the School of Life Sciences of Peking University to verify plasmid sequences. cDNAs encoding various 5-HT receptors (5-HT_1E_, 5-HT_2C_, 5-HT_5A_, and 5-HT_6_) were generated using PCR amplification of the full-length human GPCR cDNA library (hORFeome database 8.1). For optimizing the 5-HT sensor, cDNAs encoding the candidates in step 1 and step 2 were cloned into the pDisplay vector (Invitrogen) with an IgK leader sequence in the sensor upstream. In step 3, in addition to upstream IgK peptide, IRES-mCherry-CAAX cascade was fused downstream of the sensor to calibrate the membrane signal. For optimizing the linker sequence and cpGFP, site-directed mutagenesis was performed using primers containing NNB codons (48 codons, encoding 20 possible amino acids). For characterization in neurons, GRAB_5-HT1.0_ and GRAB_5-HTmut_ were cloned into the pAAV vector under the hSyn, TRE, or CAG promoter. In downstream coupling experiments, the GRAB_5-HT_ sensor and the 5-HT_2C_ receptor were cloned into the pTango and pPiggyBac vectors, respectively; two mutations were introduced into pCS7-PiggyBAC to generate hyperactive piggyBac transposase (ViewSolid Biotech)^29^. The GRAB_5-HT1.0_-SmBit and 5-HT_2C_-SmBit constructs were derived from β2AR-SmBit^13^ using a BamHI site incorporated upstream of the GGSG linker. LgBit-mGq was a generous gift from Nevin A. Lambert.

### Expression of GRAB_5-HT_ in cultured cells and *in vivo*

HEK293T cells were plated on 12-mm glass coverslips in 24-well plates and grown to 70% confluence for transfection with PEI (1 mg DNA and 3 mg PEI per well); the medium was replaced after 4-6 hours, and cells were used for imaging 24 hours after transfection. Cultured neurons were infected with AAVs expressing TRE-GRAB_5-HT1.0_ (titer: 3.8×10^13 particles/ml) and hSyn-tTA (titer: 1.3×10^14 particles/ml) or hSyn-GRAB_5-HTmut_ (titer: 1×10^13 particles/ml) at 7-9 DIV, and imaging was performed 7-14 days after infection.

For *in vivo* expression, mice were deeply anesthetized with an i.p. injection of 2,2,2-tribromoethanol (Avertin, 500 mg/kg, Sigma-Aldrich) or ketamine (10 mg/kg) and xylazine (2 mg/kg), placed in a stereotaxic frame, and the AAVs were injected using a microsyringe pump (Nanoliter 2000 Injector, WPI). For the experiments shown in Fig. 2a-e and Extended Data Fig. 4a-d, AAVs expressing CAG-GRAB_5-HT1.0_ (titer: 1.3×10^13 particles/ml) or hSyn-GRAB_5-HTmut_ (titer: 1×10^13 particles/ml) were injected (volume: 400 nl) into the DRN at the following coordinates relative to Bregma: AP: −4.3 mm, ML: 1.1 mm (depth: 2.85 mm, with a 20° ML angle). For the experiments shown in Fig. 2f,g and Extended Data Fig. 4e,f, Sindbis virus expressing GRAB_5-HT1.0_ was injected (volume: 50 nl) into the DRN at the following coordinates relative to Bregma: AP: −4.3 mm, ML: 0.0 mm (depth: 3.00 mm). For the experiments shown in Fig. 3a-d, AAVs expressing hSyn-GRAB_5-HT1.0_ (titer: 4.6×10^13 particles/ml) or hSyn-GRAB_5-HTmut_ (titer: 1×10^13 particles/ml) were injected (volume: 400 nl) into the PFC at the following coordinates relative to Bregma: AP: +2.8 mm, ML: 0.5 mm (depth: 0.5 mm). For the experiments shown in Fig. 3e-k, AAVs expressing CAG-GRAB_5-HT1.0_ or hSyn-GRAB_5-HTmut_ were injected (volume: 400 nl) into the BF at the following coordinates relative to Bregma: AP: 0 mm, ML: 1.3 mm (depth: 5.0 mm), the OFC (AP: +2.6 mm, ML: 1.7 mm, depth: 1.7 mm), and the BNST (AP: +0.14 mm, ML: 0.8 mm, depth: 3.85 mm).

### Fluorescence imaging of HEK293T cells and cultured neurons

An inverted Ti-E A1 confocal microscope (Nikon) combined with an Opera Phenix high-content screening system (PerkinElmer) were used for imaging. The confocal microscope was equipped with a 40×/1.35 NA oil-immersion objective, a 488-nm laser, and a 561-nm laser. The GFP signal was collected using a 525/50-nm emission filter, and the RFP signal was collected using a 595/50-nm emission filter. Cultured cells expressing GRAB_5-HT1.0_ or GRAB_5-HTmut_ were either bathed or perfused with Tyrode’s solution containing (in mM): 150 NaCl, 4 KCl, 2 MgCl_2_, 2 CaCl_2_, 10 HEPES, and 10 glucose (pH 7.4). Drugs were delivered via a custom-made perfusion system or via bath application. The chamber was cleaned thoroughly between experiments using 75% ethanol. Photostability was measured using confocal microscopy (1-photon illumination) with the 488-nm laser at a laser power of 350 μW, and the Opera Phenix high-content screening system was equipped with a 40×/1.1 NA water-immersion objective, a 488-nm laser, and a 561-nm laser; the GFP signal was collected using a 525/50-nm emission filter, and the RFP signal was collected with a 600/30-nm emission filter. For imaging, the culture medium was replaced with 100 μl of Tyrode’s solution, and drugs (at various concentrations) were applied in Tyrode’s solution. The fluorescence signal of the GRAB_5-HT_ sensors was calibrated using the GFP/RFP ratio.

### TANGO assay

5-HT at various concentrations was applied to 5-HT_2C_‒expressing or 5-HT1.0‒expressing HTLA cells^25^. The cells were then cultured for 12 h to allow expression of firefly luciferase (Fluc). Bright-Glo reagent (Fluc Luciferase Assay System, Promega) was then added to a final concentration of 5 μM, and luminescence was measured using a Victor X5 multilabel plate reader (PerkinElmer).

### Luciferase complementation assay

The luciferase complementation assay was performed as previously described^13^. In brief, 48 h after transfection the cells were washed with PBS, harvested by trituration, and transferred to opaque 96-well plates containing 5-HT at various concentrations. Furimazine (NanoLuc Luciferase Assay, Promega) was then added quickly to each well, followed by measurement with Nluc.

### Fluorescence imaging of GRAB_5-HT_ in brain slices

AAVs or Sindbis virus expressing GRAB_5-HT1.0_ or GRAB_5-HTmut_ were injected into the mouse DRN as described above. Three weeks after AAV injection or 18 hours after Sindbis virus injection, the mice were deeply anesthetized by an i.p. injection of Avertin or xylazine-ketamine and then transcardially perfused with 10 ml oxygenated slicing buffer consisting of (in mM): 110 choline-Cl, 2.5 KCl, 1 NaH_2_PO_4_, 25 NaHCO_3_, 7 MgCl_2_, 25 glucose, and 0.5 CaCl_2_. The mice were then decapitated, and the brains were removed and placed in cold (0-4°C) oxygenated slicing buffer for an additional 1 min. The brains were then rapidly mounted on the cutting stage of a VT1200 vibratome (Leica) for coronal sectioning at 300 μm thickness. The brain slices containing the DRN were initially allowed to recover for ≥40 min at 34°C in oxygen-saturated Ringer’s buffer consisting of (in mM): 125 NaCl, 2.5 KCl, 1 NaH_2_PO_4_, 25 NaHCO_3_, 1.3 MgCl_2_, 25 glucose, and 2 CaCl_2_. For two-photon imaging, the slices were transferred to a recording chamber that was continuously perfused with 34°C oxygen-saturated Ringer’s buffer and placed in an FV1000MPE two-photon microscope (Olympus) equipped with a 25x/1.05 NA water-immersion objective. 5-HT1.0 or 5-HTmut fluorescence was excited using a mode-locked Mai Tai Ti:Sapphire laser (Spectra-Physics) at a wavelength of 920-nm and collected via a 495-540-nm filter. For electrical stimulation, a bipolar electrode (cat. WE30031.0A3, MicroProbes) was positioned near the DRN in the slice, and imaging and stimulation were synchronized using an Arduino board with a custom-written program. The parameters of the frame scan were set to a size of 128 × 96 pixels with a speed of 0.1482 s/frame for electrical stimulation and a size of 512 × 512 pixels with a speed of 1.109 s/frame for drug perfusion experiments. For the kinetics measurements, line scans were performed with a rate of 800-850 Hz. The stimulation voltage was set at 4-6 V, and the duration of each stimulation was set at 1 ms. Drugs were bath-applied by perfusion into the recording chamber in pre-mixed Ringer’s buffer. For the Sindbis virus infected mouse brain slices, wide-field epifluorescence imaging was performed using Hamamatsu ORCA FLASH4.0 camera (Hamamatsu Photonics, Japan), and 5-HT1.0-expressing cells in acutely prepared brain slices are excited by a 460-nm ultrahigh-power low-noise LED (Prizmatix, Givat-Shmuel, Israel). The frame rate of FLASH4.0 camera was set to 10 Hz. To synchronize image capture with electrical stimulation, and fast-scan cyclic voltammetry, the camera was set to external trigger mode and triggered by a custom-written IGOR Pro 6 program (WaveMetrics, Lake Oswego, OR). For electrical stimulation, a home-made bipolar electrode was positioned near the DRN in the slice and the stimulation current was set at 350 μA and the duration of each stimulation was set at 1 ms.

### Fast-scan cyclic voltammetry (FSCV)

Carbon-fiber microelectrodes (CFME) were fabricated as described previously^30^. Briefly, cylindrical CFMEs (7 μm in radius) were constructed with T-650 carbon fiber (Cytec Engineering Materials) which was aspirated it into a glass capillary (1.2 mm O.D and 0.68 mm I.D, A-M system) and pulled using the PE-22 puller (Narishige Int.). The carbon fiber was trimmed to 50 to 70 μm in length from the pulled glass tip and sealed with epoxy which was cured at 100 °C for 2 hours followed by 150 °C overnight. CFMEs were cleaned in isopropyl alcohol for 30 min prior to the Nafion electrodeposition. Nafion was electrochemically deposited by submerging CFME tip in Nafion® solution (5 wt% 1100 EW Nafion® in methanol, Ion Power), and a constant potential of 1.0 V vs Ag/AgCl was applied to the electrode for 30 seconds. Then, Nafion coated electrodes were air dried for 10 seconds, and then at 70 °C for 10 minutes. For electrochemical detection of 5-HT, a Jackson waveform was applied to the electrode by scanning the potential from 0.2 V to 1.0 V to – 0.1 V and back to 0.2 V at 1000 V/s using a ChemClamp potentiostat (Dagan). For data collection and analysis, TarHeel (provided by R.M. Wightman, University of North Carolina) was used. For the electrode calibrations, phosphate buffer (PBS) solution was used which consisting of (in mM): 131.25 NaCl, 3.0 KCl, 10.0 NaH_2_PO_4_, 1.2 MgCl_2_, 2.0 Na2SO_4_, and 1.2 CaCl_2_ (pH 7.4). A 5-HT stock solution was prepared in 0.1 M HClO_4_ and diluted to 500 nM with PBS for calibrations prior to the experiment.

### Fluorescence imaging of transgenic flies

Fluorescence imaging in flies was performed using an Olympus two-photon microscope FV1000 equipped with a Spectra-Physics Mai Tai Ti:Sapphire laser. A 930-nm excitation laser was used for one-color imaging of 5-HT1.0 or 5-HTmut, and a 950-nm excitation laser was used for two-color imaging with 5-HT1.0 and jRCaMP1a. For detection, 495-540-nm filter for green channel and 575-630-nm filter for red channel. Adults male flies within 2 weeks post eclosion were used for imaging. To prepare the fly for imaging, adhesive tape was affixed to the head and wings. The tape above the head was excised, and the chitin head shell, air sacs, and fat bodies were carefully removed to expose the central brain. The brain was bathed continuously in adult hemolymph-like solution (AHLS) composed of (in mM): 108 NaCl, 5 KCl, 5 HEPES, 5 trehalose, 5 sucrose, 26 NaHCO_3_, 1 NaH_2_PO_4_, 2 CaCl_2_, and 1-2 MgCl_2_. For electrical stimulation, a glass electrode (resistance: 0.2 MΩ) was placed in proximity to the MB medial lobe, and voltage for stimulation was set at 10-30 V. For odorant stimulation, the odorant isoamyl acetate (cat. 306967, Sigma-Aldrich) was first diluted 200-fold in mineral oil, then diluted 5-fold with air, and delivered to the antenna at a rate of 1000 ml/min. For body shock, two copper wires were attached to the fly’s abdomen, and a 500-ms electrical stimuli was delivered at 50-80 V. For 5-HT application, the blood-brain barrier was carefully removed, and 5-HT was applied at a final concentration 100 μM. An Arduino board was used to synchronize the imaging and stimulation protocols. The sampling rate during electrical stimulation, odorant stimulation, body shock stimulation, and 5-HT perfusion was 12 Hz, 6.8 Hz, 6.8 Hz, and 1 Hz respectively.

### Two-photon imaging in mice

Fluorescence imaging in mice was performed using an Olympus two-photon microscope FV1000 equipped with a Spectra-Physics Mai Tai Ti:Sapphire laser. The excitation wavelength was 920-nm, and fluorescence was collected using a 495-540-nm filter. To perform the imaging in head-fixed mice, part of the mouse scalp was removed, and the underlying tissues and muscles were carefully removed to expose the skull. A metal recoding chamber was affixed to the skull surface with glue followed by a thin layer of dental cement to strengthen the connection. One to two days later, the skull above the prefrontal cortex was carefully removed, taking care to avoid the major blood vessels. AAVs expressing GRAB_5-HT1.0_ or GRAB_5-HTmut_ was injected as above described. A custom-made 4 mm × 4 mm square coverslip was placed over the exposed PFC and secured with glue. After surgery, mice were allowed to recover for at least three weeks. The mice were then fixed to the base and allowed to habituate for 2-3 days. During the experiment, drugs were administered by i.p. injection, and the sampling rate was 0.1 Hz.

### Fiber-photometry recording in mice

To monitor 5-HT release in various brain regions during the sleep-wake cycle, AAVs expressing GRAB_5-HT1.0_ or GRAB_5-HTmut_ were injected via a glass pipette into the BF, OFC, and BNST using a Nanoject II (Drummond Scientific). An optical fiber (200 μm, 0.37 NA) with FC ferrule was carefully inserted at the same coordinates used for virus injection. The fiber was affixed to the skull surface using dental cement. After surgery, the mice were allowed to recover for at least one week. The photometry rig was constructed using parts obtained from Doric Lens, including a fluorescence optical mini cube (FMC4_AE(405)_E(460-490)_F(500-550)_S), a blue LED (CLED_465), an LED drive (LED_2), and a photo receiver (NPM_2151_FOA_FC). To record GRAB_5-HT1.0_ and GRAB_5-HTmut_ fluorescence signals, a beam of excitation light was emitted from an LED at 20 μW, and the optical signals from GRAB_5-HT1.0_ and GRAB_5-HTmut_ were collected through optical fibers. For the fiber-photometry data, a software-controlled lock-in detection algorithm was implemented in the TDT RZ2 system using the fiber-photometry “Gizmo” in the Synapse software program (modulation frequency: 459 Hz; low-pass filter for demodulated signal: 20 Hz, 6^th^ order). The photometry data were collected with a sampling frequency of 1017 Hz. The recording fiber was bleached before recording to eliminate autofluorescence from the fiber, and the background fluorescence intensity was recorded and subtracted from the recorded signal during data analysis.

### EEG and EMG recordings

Mice were anesthetized with isoflurane (5% induction; 1.5-2% maintenance) and placed on a stereotaxic frame with a heating pad. For EEG, two stainless steel miniature screws were inserted in the skull above the visual cortex, and two additional steel screws were inserted in the skull above the frontal cortex. For EMG, two insulated EMG electrodes were inserted in the neck musculature, and a reference electrode was attached to a screw inserted in the skull above the cerebellum. The screws in the skull were affixed using thick dental cement. All experiments were performed at least one week after surgery. TDT system-3 amplifiers (RZ2 and PZ5) were used to record the EEG and EMG signals; the signal was passed through a 0.5-Hz high-pass filter at and digitized at 1526 Hz.

### Quantification and statistical analysis

Imaging data from cultured HEK293T cells, cultured neurons, acute brain slices, transgenic flies, and head-fixed mice were processed using ImageJ software (NIH) and analyzed using custom-written MATLAB programs. Traces were plotted using Origin 2018. Exponential function fitting in Origin was used to correct for slight photobleaching of the traces in Fig. 2i,m,n,o,r and Extended Data Fig. 4i,k. In Fig. 2l and Extended Data Fig. 4h, the background levels measured outside the ROI of the pseudocolor images were removed using ImageJ.

For the fiber-photometry data analysis, the raw data were binned into 1-Hz bins (i.e., down-sampled by 1000) and background autofluorescence was subtracted. For calculating ΔF/F_0_, a baseline value was obtained by fitting the autofluorescence-subtracted data with a 2^nd^ order exponential function. Slow drift was removed from the z-score‒transformed ΔF/F_0_ using the MATLAB script “BEADS” with a cut-off frequency of 0.00035 cycles/sample (https://www.mathworks.com/matlabcentral/fileexchange/49974-beads-baseline-estimation-and-denoising-with-sparsity). To quantify the change in 5-HT fluorescence across multiple animals, the z-score‒transformed ΔF/F_0_ was further normalized using the standard deviation of the signal measured during REM sleep (when there was no apparent fluctuation in the signal), yielding a normalized z-score. This normalized z-score was used for the analysis in Fig. 3e-k.

For EEG and EMG data analysis, Fast Fourier transform (FFT) was used to perform spectral analysis with a frequency resolution of 0.18 Hz. Brain state was classified semi-automatically in 5-s epochs using a MATLAB GUI and then validated manually by trained experimenters. Wakefulness was defined as desynchronized EEG activity combined with high EMG activity; NREM sleep was defined as synchronized EEG activity combined high-amplitude delta activity (0.5-4 Hz) combined with low EMG activity; and REM sleep was defined as high power at theta frequencies (6-9 Hz) combined with low EMG activity.

Except where indicated otherwise, all summary data are reported as the mean ± s.e.m. The signal to noise ratio (SNR) was calculated as the peak response divided by the standard deviation of the baseline fluorescence. Group differences were analyzed using the Student’s t-test, and differences with a p-value <0.05 were considered significant. Cartoons in Fig. 3a,e,h,j are created with BioRender.com.

### Data and software availability

The custom-written MATLAB and Arduino programs used in this study will be provided upon request.

## Acknowledgments

We thank Yi Rao for providing the two-photon microscope and Xiaoguang Lei for platform support for the Opera Phenix high-content screening system at PKU-CLS. This work was supported by the Beijing Municipal Science & Technology Commission (Z181100001318002), the Beijing Brain Initiative of Beijing Municipal Science & Technology Commission (Z181100001518004), Guangdong Grant “Key Technologies for Treatment of Brain Disorders” (2018B030332001), the General Program of National Natural Science Foundation of China (projects 31671118, 31871087, and 31925017), the NIH BRAIN Initiative (NS103558), grants from the Peking-Tsinghua Center for Life Sciences and the State Key Laboratory of Membrane Biology at Peking University School of Life Sciences (to Y.L.)., the Shanghai Municipal Science and Technology Major Project (2018SHZDZX05 to M.X.), and the Shanghai Pujiang Program (18PJ1410800 to M.X.), Alzheimer’s Association Postdoctoral Research Fellowship (P.Z.) and Peking-Tsinghua Center Excellence Postdoctoral Fellowship (Y.Z.).

## Author contributions

Y.L. conceived and supervised the project. J.W., M.J., F.D., S.H., J.F., J. Zou and S.P. performed the experiments related to developing, optimizing, and characterizing the sensor in cultured HEK293T cells and neurons. T.Q. performed the experiments using AAVs in slices. X.L. and J.Z. performed the *Drosophila* experiments. P.Z. and Y.Z. performed the experiments using the Sindbis virus in slices under the supervision of J.J.Z. J.W. performed the two-photon imaging in head-fixed mice. W.P. and K.S. performed the fiber-photometry recordings in behaving mice under the supervision of M.X. All authors contributed to the data interpretation and analysis. Y.L. and J.W. wrote the manuscript with input from all other authors.

---

**Extended Data Fig. 1:**
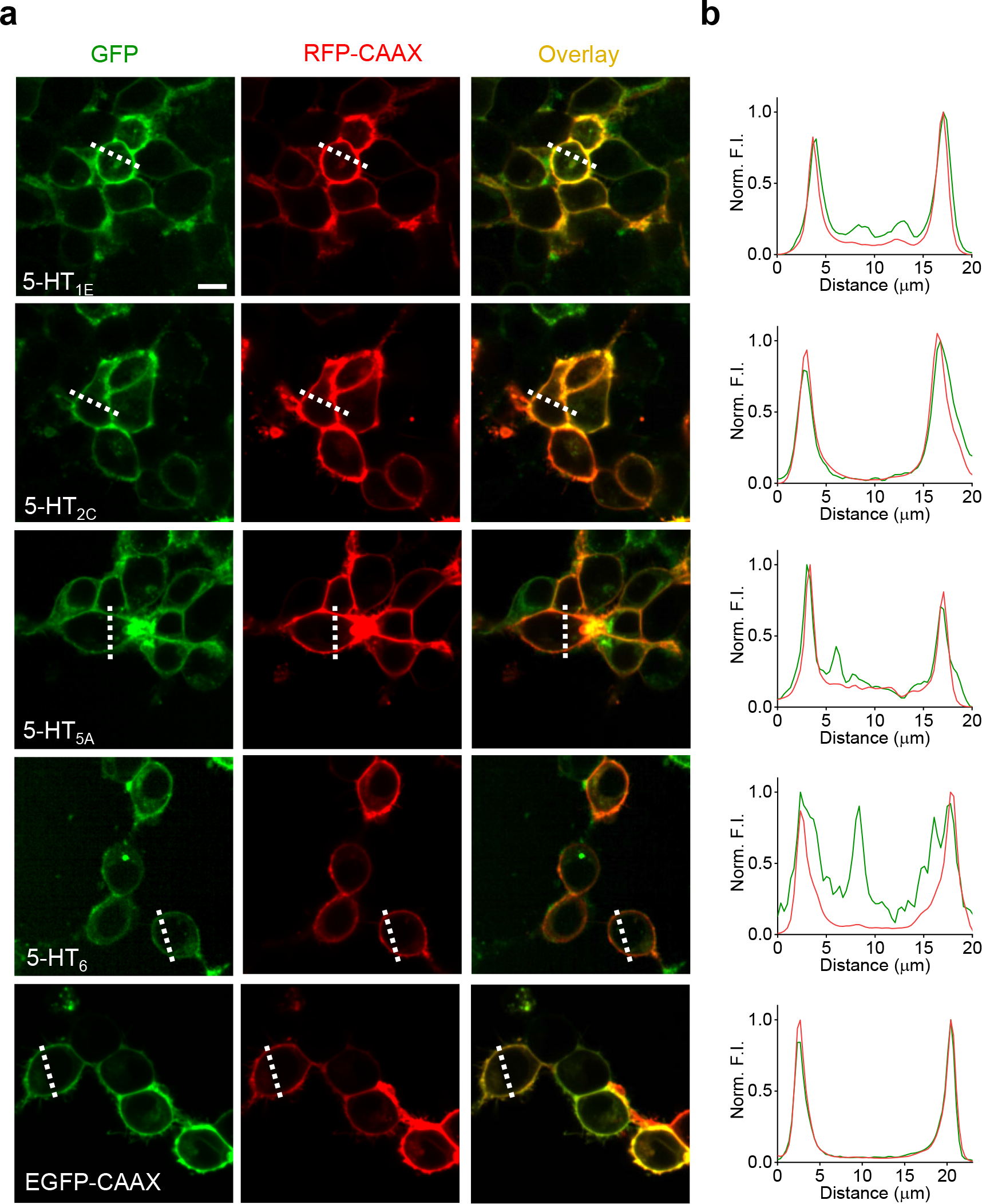
Characterization of the membrane trafficking for 5-HT receptor-based chimeras (related to Fig. 1). **(a)** Representative fluorescence images of HEK293T cells co-expressing the indicated 5-HT receptors fused with cpGFP (green) and RFP-CAAX (red); EGFP-CAAX was used as a positive control. Scale bar, 10 μm. **(b)** Normalized fluorescence intensity measured at the white dashed lines shown in **(a)** for each candidate sensor.

**Extended Data Fig. 2:**
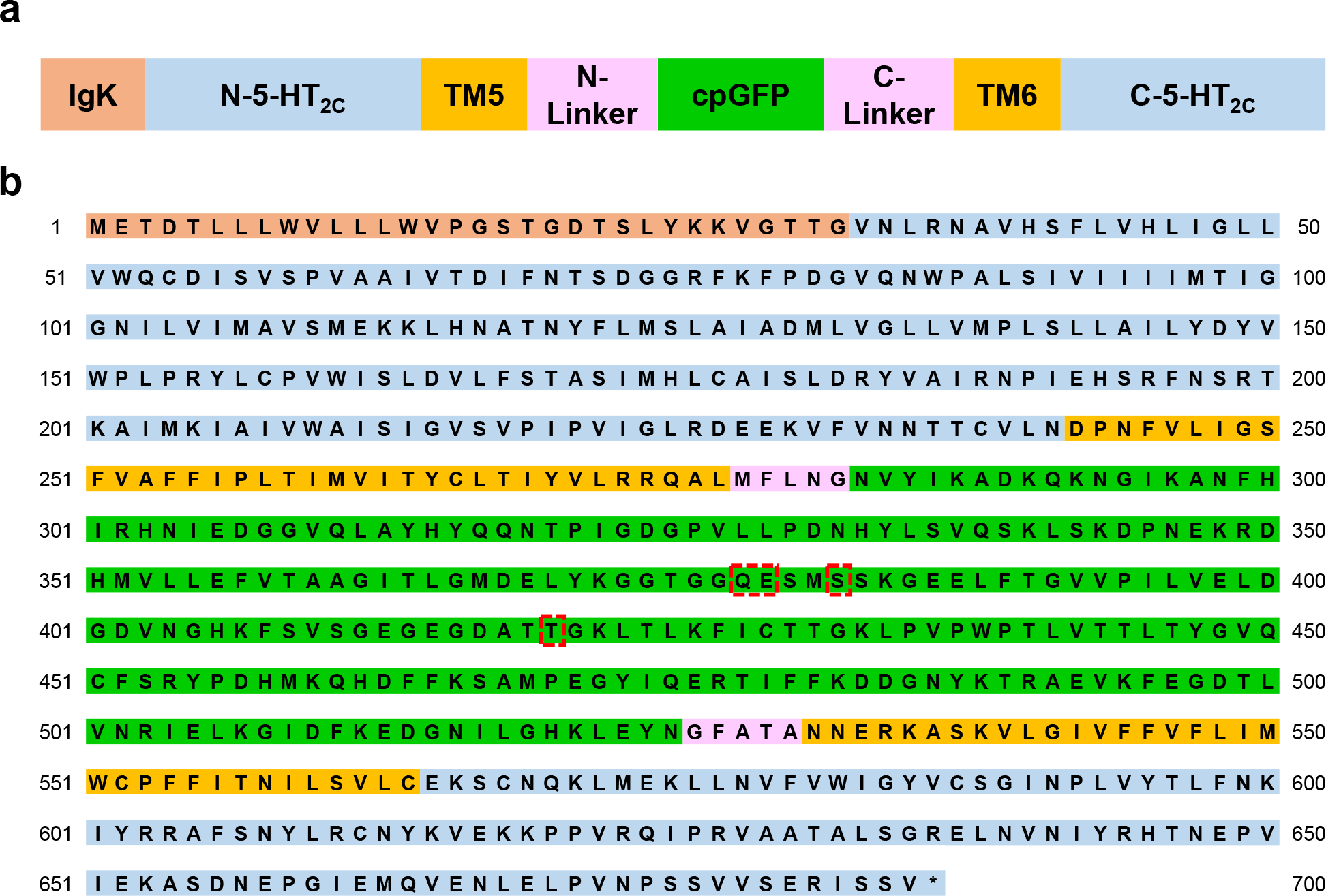
The amino acids sequence of 5-HT1.0 (related to Fig. 1). **(a)** Schematic representation of the 5-HT1.0 structure. For simplicity TM1-4, TM7 and H8 are not shown. **(b)** Amino acids sequence of the 5-HT1.0 after three steps evolution. The mutated amino acids in cpGFP (cpGFP from GCaMP6s^32^) are indicated in red box.

**Extended Data Fig. 3:**
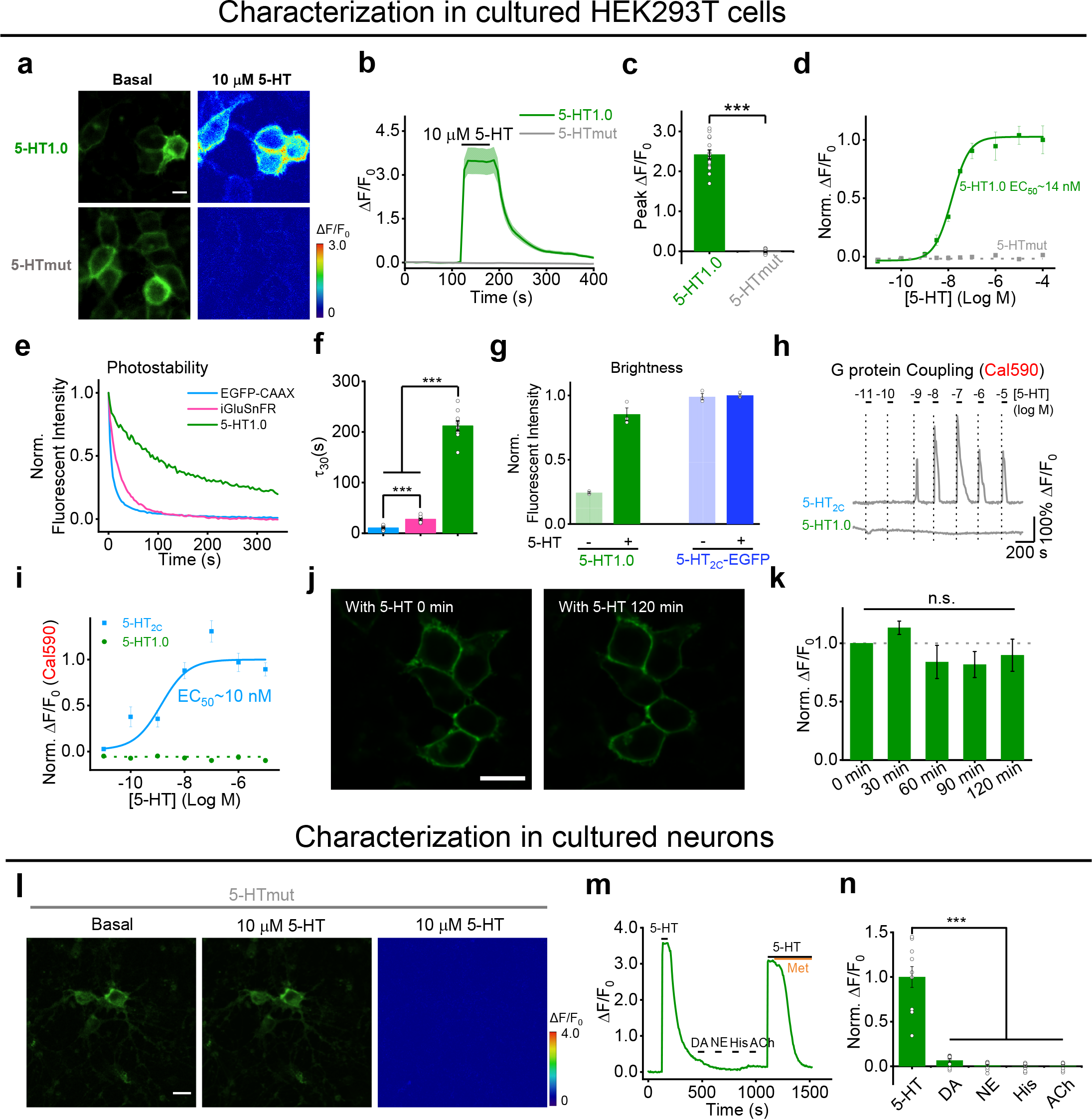
Further characterization of GRAB_5-HT_ in cultured HEK293T cells and neurons (related to Fig. 1). **(a)** Representative fluorescence and pseudocolor images of HEK293T cells expressing 5-HT1.0 or 5-HTmut before (left) and after (right) application of 10 μM 5-HT. Scale bar, 20 μm. **(b, c)** Representative fluorescence traces and group summary of the peak response in HEK293T cells expressing 5-HT1.0 or 5-HTmut; n = 14-15 cells from 3 cultures for each group. **(d)** 5-HT dose-response curves measured in cells expressing 5-HT1.0 or 5-HTmut, the EC_50_ for 5-HT1.0 is shown. **(e)** Representative normalized fluorescence measured in cells expressing 5-HT1.0, EGFP-CAAX, or iGluSnFR during continuous exposure to 488-nm laser (power: 350 μW). **(f)** Summary of the decay time constant calculated from the photobleaching curves shown in **(e)**. n = 10/3, 14/3, and 12/3 for 5-HT1.0, EGFP-CAAX, and iGluSnFR, respectively. **(g)** Summary of the brightness measured in cells expressing 5-HT1.0 or 5-HT_2C_-EGFP in the absence or presence of 10 μM 5-HT, normalized to the 5-HT_2C_-EGFP + 5-HT group; n = 3 wells per group with 300–500 cells per well. **(h, i)** Intracellular Ca^2+^ measured in cells expressing 5-HT1.0 or the 5-HT2C receptor and loaded with the red fluorescent calcium dye Cal590. Representative traces are shown in **(h)**, and the peak responses are plotted against 5-HT concentration in **(i)**; n = 15/3 for each group. **(j, k)** Fluorescence response of 5-HT1.0 expressing cells to 5-HT perfusion for two hours. Representative fluorescence images **(j)** and the summary data **(k)** showing the response to 10 μM 5-HT applied at 30 min intervals to cells expressing 5-HT1.0; n = 3 wells per group with 100-300 cells per well. Scale bar, 20 μm. **(l)** Cultured cortical neurons expressing the 5-HTmut sensor were imaged before (left) and after (middle) 5-HT application. The pseudocolor image at the right shows the change in fluorescence. Scale bar, 20 μm. **(m, n)** Representative trace **(m)** and group summary **(n)** of cultured neurons expressing 5-HT1.0 in response to indicated compounds at 10 μM each; in **(m)**, Met was applied where indicated; n = 9/3.

**Extended Data Fig. 4:**
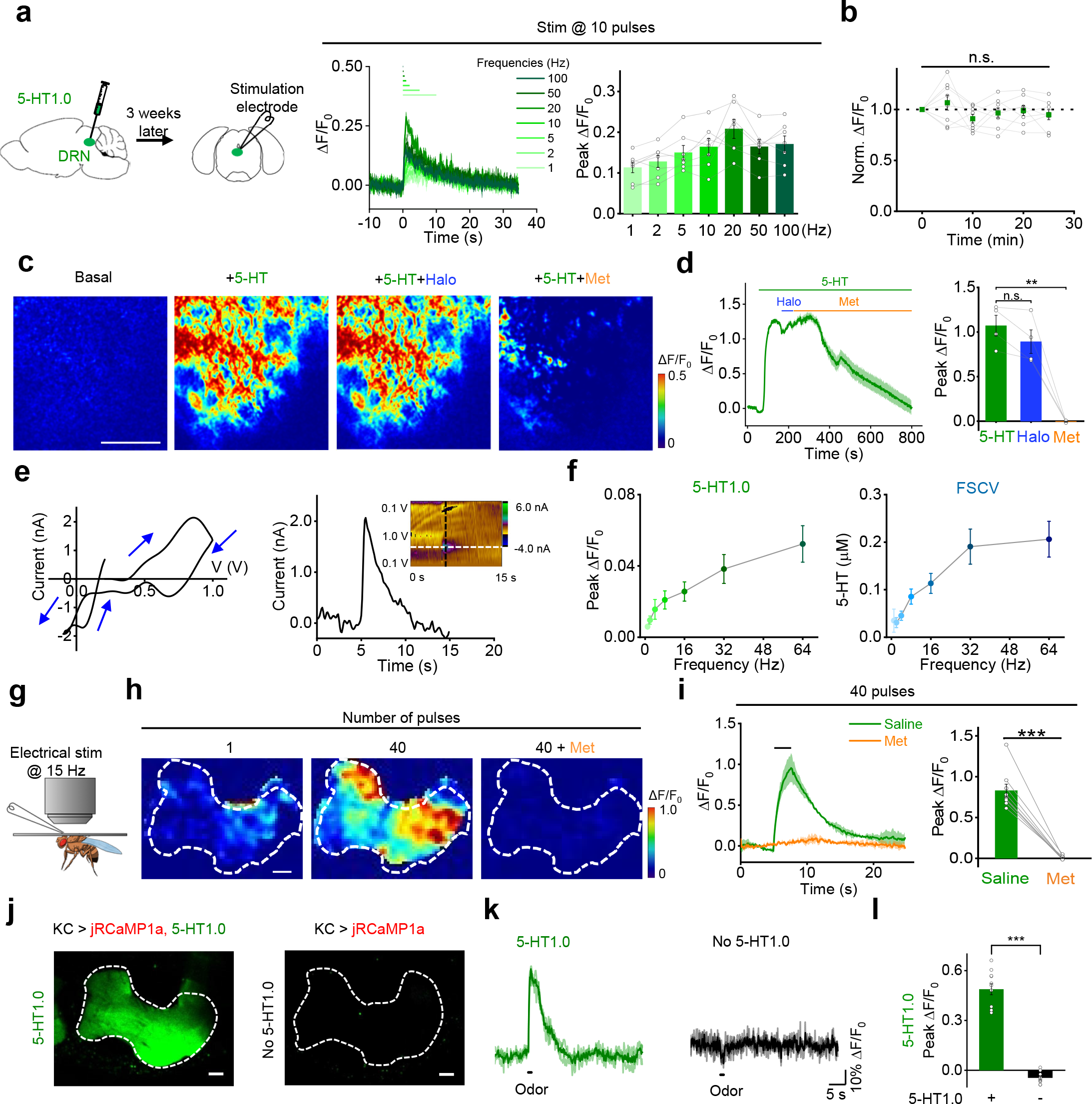
Probing endogenous 5-HT release in mouse brain slices and *Drosophila in vivo* (related to Fig. 2). **(a)** Left: schematic diagram depicting the acute mouse brain slice preparation, with AAV-mediated expression of 5-HT1.0 in the DRN. Middle and right: fluorescence traces (middle) and group data (right) of the change in 5-HT1.0 fluorescence in response to 10 electrical stimuli applied at the indicated frequencies; n = 7 slices from 5 mice. **(b)** Summary of the change in 5-HT1.0 fluorescence in response to 6 trains of electrical stimuli (20 pulses at 20 Hz) delivered at 5-min intervals. The responses are normalized to the first train; n = 8 slices from 5 mice. **(c, d)** Representative pseudocolor images **(c)**, fluorescence traces **(d, left)**, and group data **(d, right)** of 5-HT1.0 fluorescence in response to perfusion of 5-HT, 5-HT+Halo, and 5-HT+Met; n = 4 slices from 3 mice for each group. **(e)** Left: representative FSCV data of 5-HT release in DRN. A specific 5-HT waveform (0.2 V to 1.0 V and ramped down to – 0.1 V, and back to 0.2 V at a scan rate of 1000 V/s) was applied to the CFME at a frequency of 10 Hz. Right: current vs time traces is extracted at horizontal white dashed line shows immediate increase in 5-HT response after electrical stimulation (20 pulses, 2 ms pulse width, 64 Hz). A cyclic voltammogram (inset) is extracted at the vertical black dashed line shows oxidation and reduction peaks at 0.8 V and 0 V, respectively. **(f)** Left: group data of fluorescence response in 5-HT1.0-expressing DRN neurons to electrical stimuli with varied frequency delivered at 20 pulses. Right: average data of peak 5-HT concentration measured by FSCV at varied stimulating frequencies; n = 11 neurons from 9 mice. **(g)** Schematic drawing showing *in vivo* two-photon imaging of a *Drosophila*, with the stimulating electrode positioned near the mushroom body (MB). **(h, i)** Representative pseudocolor images **(h)**, fluorescence traces and group summary **(i)** of the change in 5-HT1.0 fluorescence in the MB horizontal lobe in response to 40 electrical stimuli at 15 Hz in control (saline) or 10 μM Met; n = 9 flies for each group. Scale bar, 10μm. **(j)** Fluorescence images of green channel of MB in flies with co-expression of jRCaMP1a and 5-HT1.0 (left) or expressing jRCaMP1a alone (right) in the KCs. Scale bar, 10 μm. **(k, l)** Representative traces **(k)** and group summary **(l)** of 5-HT signal in the MB of flies co-expressing jRCaMP1a and 5-HT1.0 or jRCaMP1a alone; where indicated, a 1-s odorant stimulation was applied; n = 10-11 flies for each group.

**Extended Data video 1.**
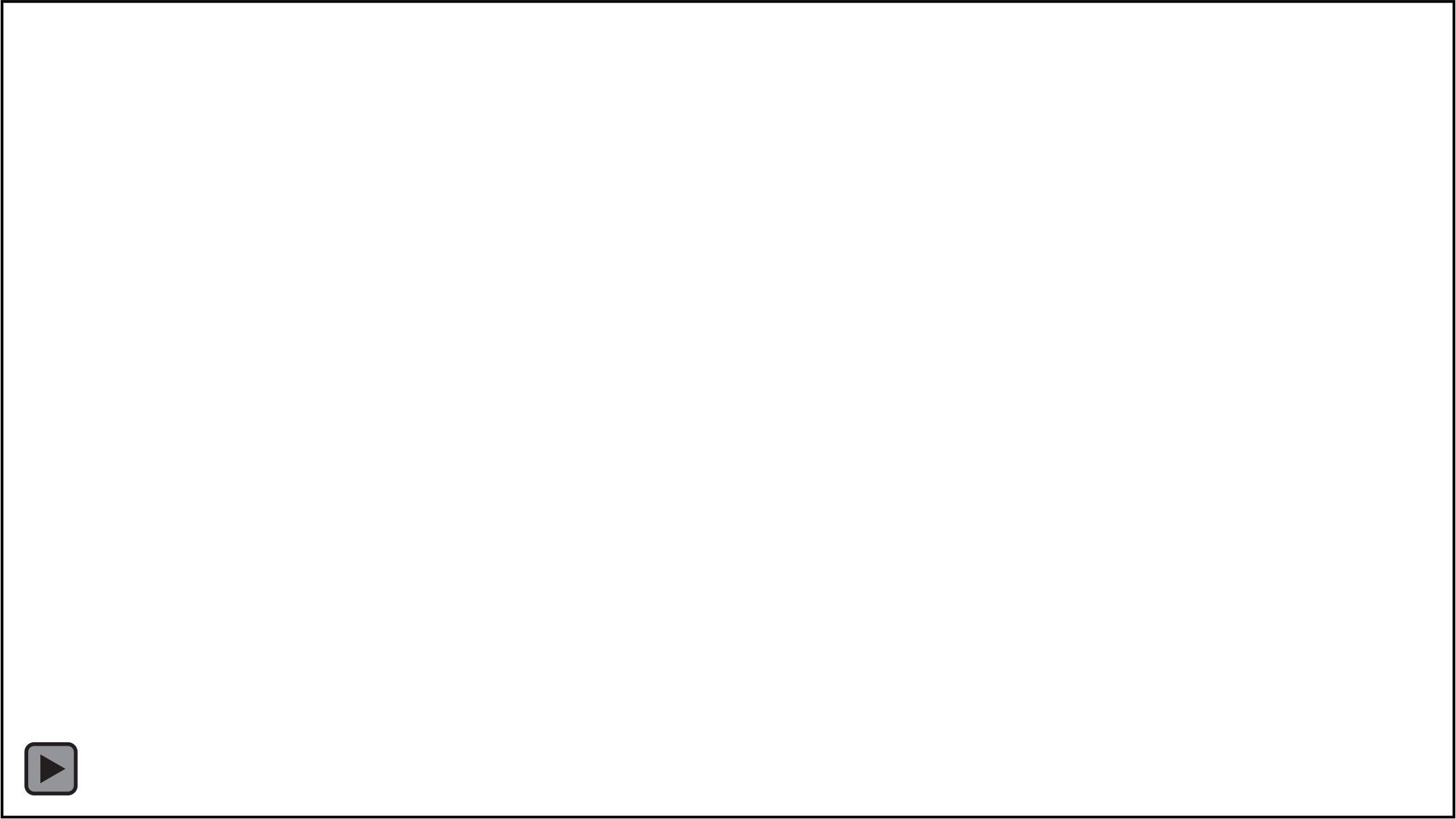
GRAB_5-HT_ reports the sensory-relevant 5-HT release in *Drosophila* (Related to Fig. 3k-p). Fluorescence responses of 5-HT1.0 and 5-HTmut in the MB β’ lobe measured in response to a 1-s odor application, a 0.5-s body shock, and application of 100 μM 5-HT.

